# Gut microbially produced phenolic acids have differential and sex-dependent effects on host metabolic health

**DOI:** 10.64898/2026.10.12.757096

**Authors:** Jiyeon Kim, Kelley M. Carr, Sara Alqudah, Kathryn A. Bacik, Hannah Park, Clifton G. Fulmer, Muyiwa Awoniyi, Jan Claesen, Ina Nemet

**Author notes:** Corresponding author: Ina Nemet, 9500 Euclid Ave, NC1-136, Cleveland Clinic, Cleveland, OH, USA 44195. Current address: Department of Pathology, University of Chicago, Chicago, IL, USA. Current address: Department of Engineering Science and Mechanics, Penn State College of Engineering, University Park, PA, USA.

## Abstract

Gut microbiota-derived metabolites are increasingly recognized as important regulators of host metabolism, but the specific functions of individual metabolites remain poorly understood. We therefore investigated the metabolic effects of two structurally related, phenylalanine-derived microbial metabolites, phenylacetic acid (PAA) and phenylpropionic acid (PPA), in a mouse model of diet-induced obesity.

Although clinical studies have linked PAA and its downstream product phenylacetylglutamine to metabolic disease, PAA did not alter body mass or adiposity in this model. Instead, it improved glucose tolerance selectively in male mice and markedly remodeled lipid metabolism in both sexes, reducing circulating triglycerides and acylcarnitines and activating hepatic PPAR-α-associated transcriptional programs associated with fatty acid ω-hydroxylation and peroxisomal lipid metabolism. Despite this extensive metabolic remodeling, PAA did not worsen hepatic steatosis, fibrosis, or markers of liver injury.

By contrast, PPA had minimal effects on hepatic metabolic pathways but produced pronounced sex-dependent physiological responses, increasing adiposity in female mice and improving glucose tolerance in male mice.

These findings show that structurally related microbial metabolites can have distinct, sex-dependent metabolic effects and highlight the need to define the actions of individual microbiome-derived metabolites when developing targeted therapies for metabolic disease.

**Graphical Abstract:** 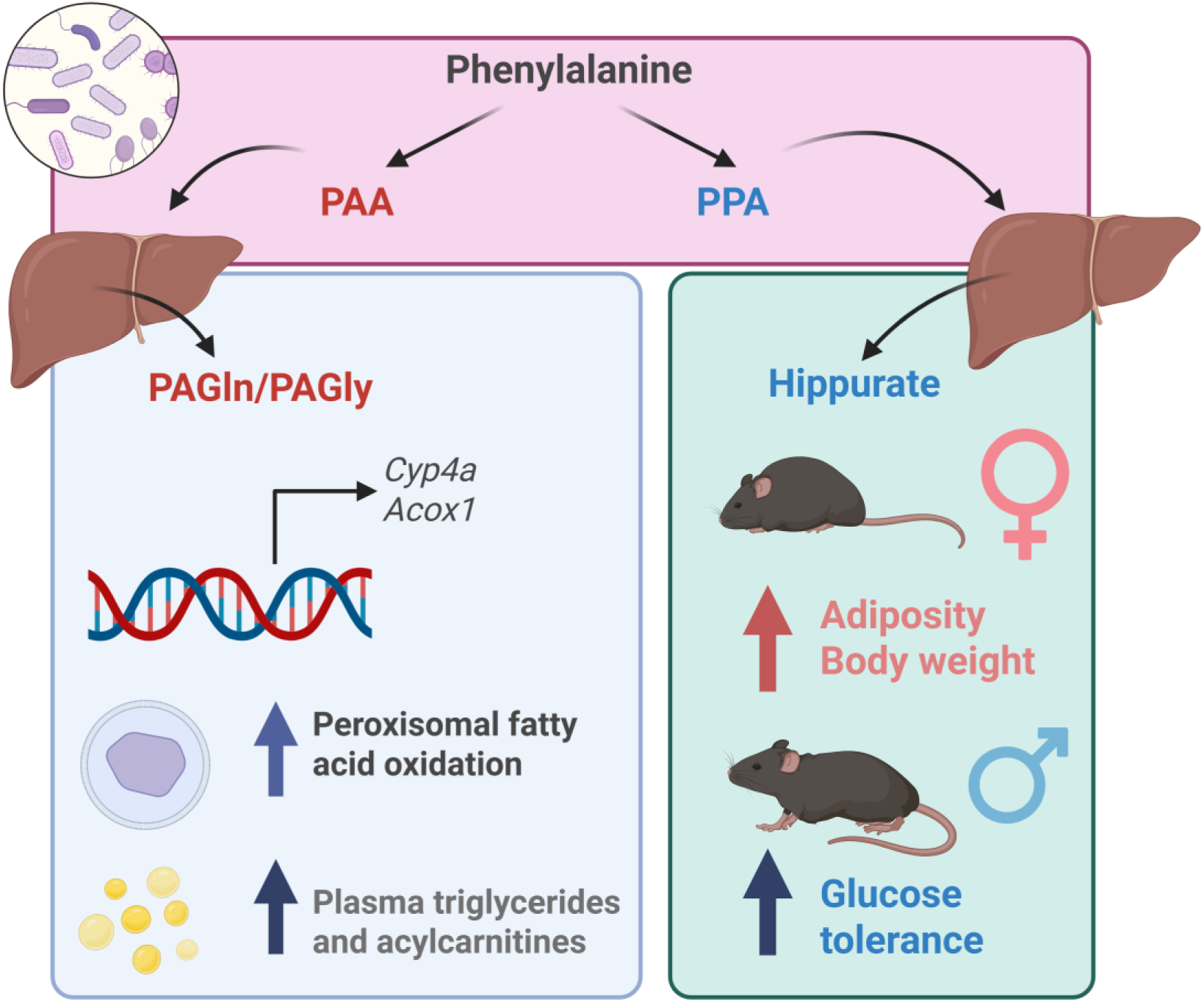

## 1. Introduction

Large clinical cohort studies have consistently reported alterations in gut microbial composition in individuals with obesity, metabolic syndrome, and type 2 diabetes mellitus (T2DM), supporting a link between the gut microbiome and metabolic disease [1–5]. More recently, analyses of ethnically and geographically diverse populations have further demonstrated that gut microbiome composition is associated with the future risk of developing T2DM [6]. Causal evidence from preclinical models reinforces these observations, as fecal microbiota transplantation from glucose-intolerant mice into germ-free recipients induces glucose intolerance [7], whereas, in humans, transfer of microbiota from lean donors to individuals with metabolic syndrome improves microbial diversity and insulin sensitivity [8]. Although these findings implicate gut microbial dysbiosis in metabolic dysfunction, the underlying mechanisms remain incompletely understood.

One major pathway through which the gut microbiota influences host physiology is the production of bioactive metabolites that exert both beneficial and deleterious effects on metabolic health. For example, dietary fibers and resistant starches are fermented by gut bacteria into short-chain fatty acids (SCFAs), including acetate, propionate, and butyrate, which regulate energy balance, insulin sensitivity, gut homeostasis, and blood pressure through G protein–coupled receptor signaling and epigenetic mechanisms such as histone deacetylase inhibition [9–15]. Similarly, microbial conversion of primary bile acids synthesized by the host into secondary bile acids modulates lipid and glucose metabolism through signaling via farnesoid X receptor (FXR) and Takeda G protein-coupled receptor 5 (TGR5) [16–19]. These well-established pathways illustrate how microbial metabolites serve as intermediates linking diet, microbiome function, and host metabolic regulation. In addition to these pathways, microbial metabolism of dietary amino acids has emerged as an important and less well-characterized contributor to host metabolic homeostasis. Dietary phenylalanine that escapes host absorption is metabolized by gut bacteria into phenylpyruvic acid, which is subsequently converted through oxidative or reductive pathways to phenylacetic acid (PAA) or phenylpropionic acid (PPA), respectively [20]. Following absorption into the portal circulation, these microbial intermediates undergo further host hepatic metabolism. PAA is conjugated with glutamine in humans to form phenylacetylglutamine (PAGln) and with glycine in rodents to form phenylacetylglycine (PAGly) [21, 22], whereas PPA is first metabolized via β-oxidation and subsequently conjugated with glycine to generate benzoylglycine commonly known as hippuric acid or hippurate [23] (**Figure 1A**). These host–microbial co-metabolites represent circulating signals that may link microbial activity to systemic metabolic regulation.

**Figure 1.**
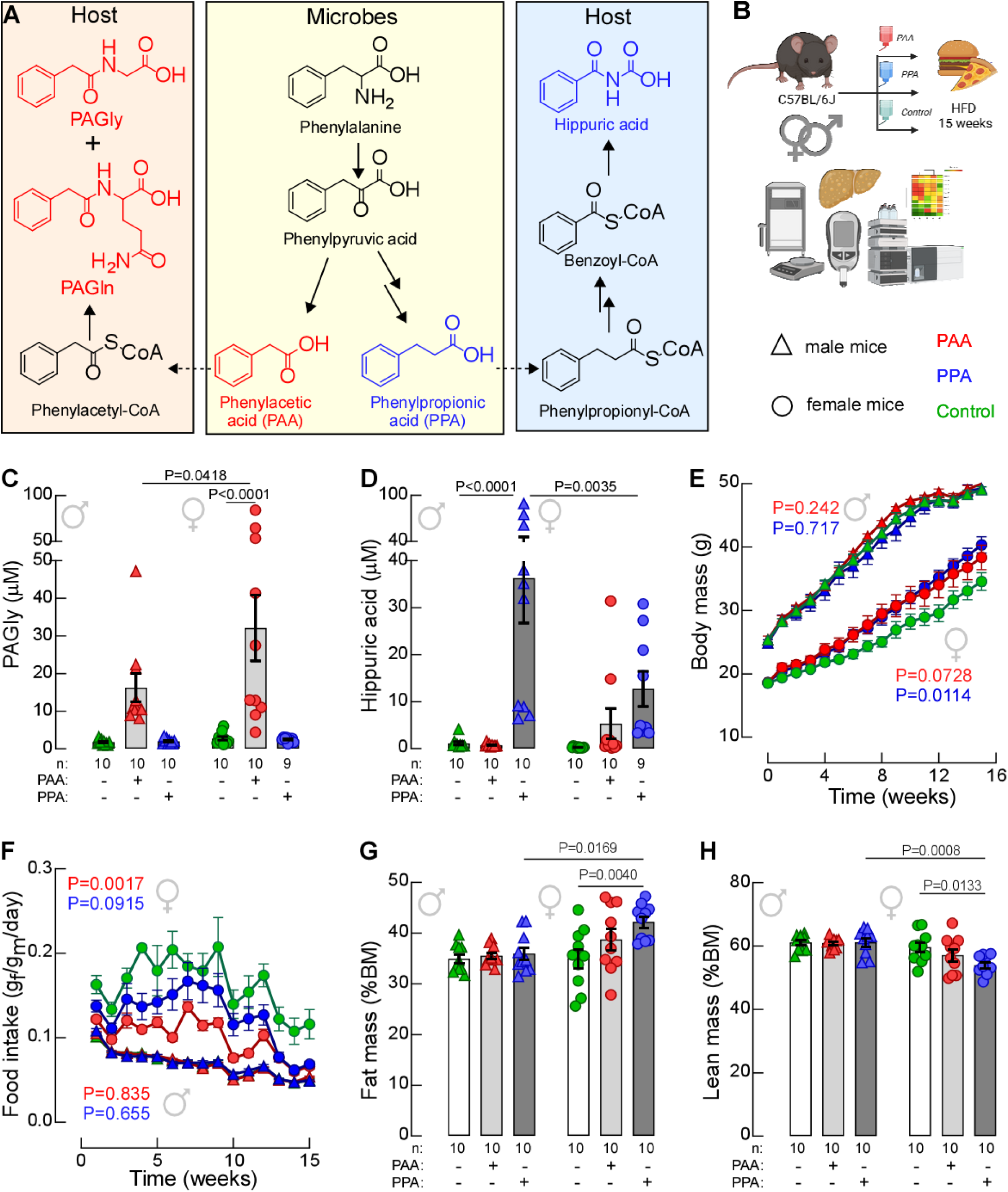
Phenylpropionic acid (PPA) promoted higher body mass and fat gain in female C57BL/6J mice on a high-fat diet. (**A**) Metaorganismal production of phenylacetylglutamine (PAGln), phenylacetylglycine (PAGly) and hippuric acid from respective microbially produced PAA and PPA. (**B**) Experimental design. Circulating levels of (**C**) PAGly and (**D**) hippuric acid. (**E**) Body mass over time, (**F**) food intake, and (**G**) body fat and (**H**) lean mass expressed as percentage of body mass (%BM) at the end of the study in male (triangles) and female (circles) mice on a high-fat diet (HFD) receiving dietary PAA (red) or PPA (blue) supplementation or a control diet (green). Data were analyzed by two-way ANOVA followed by Šídák’s multiple comparisons test (**E** and **F**: red *P*-values indicate significant differences between control and PAA, whereas blue *P*-values indicate significant differences between control and PPA). Values are mean ± SEM; n = 9-10 as indicated.

Clinical data suggest that these phenylalanine-derived metabolites exhibit divergent associations with metabolic diseases. Elevated circulating PAGln levels have been associated with T2DM [21, 24, 25] and other conditions linked to increased metabolic risk, including human immunodeficiency virus (HIV) infection [26] and alcohol use disorders [27]. In obese women without T2DM, circulating levels of PAA are positively associated with hepatic steatosis and inversely associated with microbial gene richness [28]. In contrast, hippuric acid has been consistently associated with favorable metabolic phenotypes, including a negative association with metabolic dysfunction-associated steatotic liver disease (MASLD) [28, 29], increased microbial diversity, higher intake of fiber-rich foods, lower prevalence of metabolic syndrome, and greater physical activity [30–32]. Consistent with these observations, circulating and urinary hippuric acid levels are reduced in obesity and T2DM, although some smaller studies report conflicting findings [33, 34], underscoring the complexity of these associations.

Several preliminary experimental studies support the concept that these metabolites exert functionally distinct effects on host metabolism. *In vitro*, PAA promotes lipid accumulation and alters the expression of genes involved in glucose and lipid metabolism in primary human hepatocytes [28], whereas hippuric acid produces the opposite effect in immortalized human hepatocytes [29]. *In vivo*, short-term exposure to PAA increases hepatic triglyceride accumulation in mice [28], whereas chronic hippuric acid administration improves glucose tolerance in male mice fed a high-fat diet [31].

Despite these insights, the pathways through which these metabolites influence systemic metabolism, and the extent to which their effects are context- and sex-dependent, remain poorly defined. Here, using a mouse model of diet-induced obesity, we demonstrate that dietary PAA remodels the circulating lipidome and alters hepatic transcriptional programs involved in lipid metabolism and homeostasis without significantly affecting adiposity or steatosis progression. In contrast, supplementation with PPA, the microbial precursor of hippuric acid, increases body mass and adiposity in female mice while improving glucose tolerance in male mice.

These findings redefine phenylalanine-derived microbial metabolism as a context- and sex-dependent regulator of host metabolic physiology and establish a previously unrecognized axis through which the gut microbiome shapes systemic metabolic homeostasis.

## 2. Materials and methods

### 2.1. Experimental design and mice

All animal studies were approved by the Cleveland Clinic Research Institutional Animal Care and Use Committee. Male and female C57BL/6J mice were obtained from Jackson Laboratories and housed under a standard 12-hour light/dark cycle in the Cleveland Clinic Lerner Research Building mouse facility. Mice were fed a high-fat diet (Research Diets, Cat# D12492i) consisting of 31.7% fat (60.0% kcal from fat). PAA and PPA (1% w/v; Sigma-Aldrich, Cat# W287806 and Cat# W288918, respectively) were administered in drinking water. Both PAA and PPA were neutralized with sodium bicarbonate (Sigma-Aldrich, Cat# S6297) and supplemented with an equimolar amount of glycine (Sigma-Aldrich, Cat# G7126) to avoid glycine depletion resulting from PAGly and hippuric acid formation. Control mice received an equivalent amount of glycine as a sodium salt (sodium glycinate; Sigma-Aldrich, Cat# G6761). Non-fasting blood was collected from the saphenous vein between 7.00 and 9.00 a.m. using heparin-coated capillaries. Blood was centrifuged (1,500 x *g*, 10 min, 4 °C) and plasma was aliquoted and stored at −80 °C until analysis. At the end of the study, mice were fasted for 4 h and euthanized by anesthesia overdose (>300 mg/kg ketamine + 30 mg/kg xylazine). Blood was collected by cardiac puncture. Tissues were harvested, immediately flash-frozen in liquid nitrogen, and stored at −80 °C until analysis.

### 2.2. Phenotyping of mouse models

Body mass and food intake were measured at baseline and weekly throughout the study. Lean and fat body mass were assessed using EchoMRI body composition analysis (EchoMRI, Houston, TX). Blood glucose was measured using a handheld Accu-Chek Guide glucose meter (Roche Diagnostics Operations, Indianapolis, IN). Plasma insulin was measured using an Ultra-Sensitive Mouse Insulin ELISA kit (Crystal Chem, Cat# 90080) according to the manufacturer’s instructions. Homeostatic Model Assessment for Insulin Resistance (HOMA-IR) was calculated as follows: [insulin (µIU/mL) x blood glucose (mmol/L)]/22.5.

Intraperitoneal (IP) glucose tolerance tests (GTTs) and insulin tolerance tests (ITTs) were performed in mice fasted for 4 h. Mice received a single IP injection of glucose (1.0 g/kg body mass) or insulin (0.5 U/kg body mass), respectively. Blood glucose was measured in duplicate before injection (0 min) and at 15, 30, 60, and 120 min after injection.

Total plasma free fatty acids were measured using enzymatic colorimetric assays (HR Series NEFA-HR, Wako, Cat# 999-34691) according to the manufacturer’s instructions.

### 2.3. Histological analysis

Hematoxylin and Eosin (H&E) staining was performed on a Leica Spectra automated stainer. Sections were deparaffinized in xylene, rehydrated through graded ethanol, stained with hematoxylin, subjected to acid differentiation and bluing, counterstained with eosin Y, dehydrated, and cleared in xylene. H&E-stained liver sections were evaluated by a pathologist blinded to treatment groups for hepatic steatosis, lobular inflammation, and hepatocyte ballooning, and the nonalcoholic fatty liver disease activity score (NAS) was calculated as the sum of these three components.

For collagen content quantification, sections were deparaffinized in xylene and hydrated in distilled water. Nuclei were stained with Weigert’s hematoxylin for 8 min, followed by incubation in Picrosirius red solution for 60 min. Slides were rinsed twice in acidified water (0.5% v/v glacial acetic acid in distilled water), dehydrated in two changes of absolute ethanol, cleared, and coverslipped. Brightfield scans of the Picrosirius red-stained tissue sections were captured at 20× magnification (0.5 µm/pixel resolution) using an Aperio AT2 whole-slide scanning system (Leica Biosystems GmbH, Germany). Collagen area as a percentage of total tissue area was measured using QuPath [35]. Sirius red and picric acid colors were separated using their color vectors and color deconvolution in QuPath. A threshold-based classifier was used to detect and annotate liver sections, followed by manual curation of the annotations to remove areas with artifacts. An artificial neural network (ANN)-based pixel classifier was used to detect collagen in Picrosirius red-stained liver sections. The ANN-based pixel classifier was trained on 100 patches of 512 × 512 pixels to detect collagen. These training regions were extracted from across the image dataset and represented heterogeneous Sirius red staining. Total tissue area and collagen area were reported for two serial liver sections per mouse.

### 2.4. Plasma ALT, AST, and ALP activity

Plasma alanine aminotransferase (ALT), aspartate aminotransferase (AST), and alkaline phosphatase (ALP) activities were measured using enzymatic assays according to the manufacturers’ instructions. ALT and AST were measured using Sekure Chemistry kits (Cat# 318-30 and Cat# 319-30, respectively) and ALP was measured using a Novus Biologicals kit (Cat# NBP3-24466).

### 2.5. Quantification of hepatic lipids

Liver lipids were extracted and total triglycerides, free cholesterol, and total cholesterol were quantified using enzymatic colorimetric assays as previously described [36]. Briefly, liver samples were extracted in chloroform: methanol (2:1 v/v, 3 mL) at 55 °C for 2 h, followed by overnight incubation at room temperature. An aliquot (2 mL) of the extract was transferred to a solution of Triton X-100 in chloroform (1%, 2 mL), vortexed, and dried under a constant stream of nitrogen gas. The dry residue was resuspended in Milli-Q water (500 µL) before quantification of free cholesterol (Free Cholesterol E, Wako, Cat# 999-02501), total cholesterol (Infinity Cholesterol liquid stable reagent, Thermo Scientific, Cat # TR13421), and triglycerides (L-Type Triglyceride M, Wako, Cat# 999-02891) according to the manufacturer’s instructions. Standards and blanks were processed in the same manner as samples, and final values were normalized to liver mass.

### 2.6. Targeted liquid chromatography-tandem mass spectrometry (LC-MS/MS) analysis

All targeted LC-MS/MS analyses were performed using a chromatographic system consisting of two Shimadzu LC-30AD pumps (Nexera X2), a CTO-20AC oven operating at 40 °C, and a SIL-30AC-MP autosampler coupled to an 8050 triple-quadrupole mass spectrometer (Shimadzu Scientific Instruments, Inc.). The following ion source parameters were applied: nebulizing gas flow, 3 L/min; heating gas flow, 10 L/min; interface temperature, 300 °C; desolvation line temperature, 250 °C; heat block temperature, 400 °C; and drying gas flow, 10 L/min. Data were analyzed using LabSolutions software version 5.89 (Shimadzu).

#### 2.6.1. PAA, PAGln, PAGly, PPA, and hippuric acid quantification

PAA, PAGln, PAGly, PPA, and hippuric acid were quantified using a previously published stable-isotope-dilution LC-MS/MS method [37], with some modifications. Briefly, ice-cold methanolic internal standard solution containing D5-PAGln, D5-PAGly, D5-PAA, D5-hippuric acid, and D9-PPA (80 µL) was added to plasma samples (20 µL), followed by vortexing and centrifuging (21,000 x *g*; 4 °C; 15 min). The clear supernatants were transferred to glass vials with micro-inserts. Chromatographic separation was performed using an XSelect HSS T3 column (3.0 x 100 mm; 3.5 µm; Waters, Ireland, Cat# 186004780). A gradient of solvent A (0.1% acetic acid in water) and B (0.1% acetic acid in acetonitrile) was used at a flow rate of 0.4 mL/min with a 1 µL injection volume. Electrospray ionization was performed in both positive and negative ion modes using multiple reaction monitoring (MRM). The following transitions were monitored in positive ion mode: *m/z* 265.2 ◊ 130.1 for PAGln; *m/z* 271.0 ◊ 130.1 for D5-PAGln; *m/z* 193.8 → 76.1 for PAGly; *m/z* 198.9 ◊ 76.1 for D5-PAGly; 180.0 → 105.1 for hippuric acid; and *m/z* 185.0 ◊ 110.1 for D5-hippuric acid. The following transitions were monitored in negative ion mode: *m/z* 135.0 → 91.0 for PAA; *m/z* 140.2 ◊ 95.9 for D5-PAA; *m/z* 149.0 → 105.1 for PPA; *m/z* 158.2 ◊ 114.1 for D9-PPA.

#### 2.6.2. Bile acids

Bile acids were quantified by a stable-isotope-dilution LC-MS/MS method as previously described [38]. Briefly, ice-cold methanolic internal standard solution (80 µL) containing D4-chenodeoxycholic acid, D4-cholic acid, D4-deoxycholic acid, D4-lithocholic acid, D4-glycochenodeoxycholic acid, D4-glycocholic acid, D4-glycodeoxycholic acid, D4-glycolithocholic acid, D4-glycoursodeoxycholic acid, D4-taurocholic acid, D4-taurodeoxycholic acid, and D4-taurochenodeoxycholic acid was mixed with liver methanolic extract (20 µL), vortexed and centrifuged (14,000 x *g*, 20 min, 4 °C). The supernatant was transferred to glass vials with micro-inserts. Chromatographic separation was performed on an ACE Excel C18-Amide column (75 mm × 2.1 mm; 1.7 μm; Avantor, Cat# EXL-1712-7502U). A gradient of solvent A (0.1% acetic acid in water) and B (0.1% acetic acid in acetonitrile: methanol 50:50; v/v) was used for chromatographic separation at a flow rate of 0.4 mL/min and a 3 µL injection volume.

#### 2.6.3. Acylcarnitines

Acylcarnitines were quantified in mouse plasma, feces, heart, skeletal muscle, and liver using stable-isotope-dilution LC-MS/MS, as previously described [39]. Briefly, ice-cold methanolic internal standard solution (80 µL) containing D3-L-carnitine (D3-Car 0:0), acetyl-D3-L-carnitine (D3-Car 2:0), butyryl-D7-L-carnitine (D7-Car 4:0), hexanoyl-D3-L-carnitine (D3-Car 6:0), and hexadecanoyl-D3-L-carnitine (D3-Car 16:0) was added to mouse plasma, fecal or tissue methanolic extracts (20 µL), followed by vortexing and centrifugation (14,000 x *g*, 15 min, 4 °C). The supernatant was transferred to glass vials with microinserts and subjected to LC-MS/MS analysis. Chromatographic separation was achieved using a Kinetex HILIC column (100 x 2.1; 2.6 μm; Pheno mm, Cat# PRD-509459). A gradient of 10 mM ammonium formate and 0.1% formic acid in water (solvent A) and 10 mM ammonium formate and 0.1% formic acid in 95% acetonitrile (solvent B) was used for chromatographic separation at a flow rate of 0.4 mL/min with a 1 µL injection volume.

Short-chain (SC) acylcarnitines were calculated as the sum of individual C2–C5 acylcarnitines, medium-chain (MC) acylcarnitines as the sum of C6–C12 acylcarnitines, and long-chain (LC) acylcarnitines as the sum of C14–C18 acylcarnitines.

### 2.7. Untargeted lipidomics

Untargeted lipidomic analysis of plasma samples from mice fasted for 4 h was performed by MS-Omics (Vedbaek, Denmark) using an Elute-XT LC system (Bruker Daltonics, Germany) coupled to a trapped ion-mobility high-resolution quadrupole-time-of-flight mass spectrometer (timsTOF Pro 2; Bruker Daltonics, Germany) equipped with a vacuum-insulated probe-heated electrospray ionization source. Ionization was performed in positive and negative ionization modes using an electrospray ionization interface. Lipids were separated on a Waters® ACQUITY Charged Surface Hybrid (CSH™) C18 column (2.1 x 100 mm, 1.7 µm, Cat# 186002352) maintained at 65 °C. Mobile phase A consisted of acetonitrile/water (60:40, v/v) and mobile phase B consisted of isopropanol/acetonitrile (90:10; v/v); both mobile phases contained 10 mM ammonium formate and 0.1% formic acid. Lipids were eluted using a gradient in which mobile phase B was increased from 40% to 99% over 4 min with a flow rate of 0.55 mL/min. Peak areas were extracted using MetaboScape 2023 software (Bruker Daltonics, Germany).

Compound identifications were assigned at four confidence levels: Level 1, identification based on retention times compared with in-house authentic standards, accurate mass within 3 ppm, and tandem mass spectrometry (MS/MS) spectra; Level 2a, identification based on retention time compared with in-house authentic standards and accurate mass within 3 ppm; Level 2b, identification based on accurate mass within 3 ppm and MS/MS spectra; and Level 3, identification based on accurate mass within 3 ppm alone. Lipids identified at Levels 1 and 2 were used for the analysis.

### 2.8. Cell culture

Human hepatoma (HuH-7) cells were seeded in 6-well tissue culture plates (2×10^5^ cells/well; Corning, Cat# 3506) and cultured in Dulbecco’s Modified Eagle Medium supplemented with 10% fetal bovine serum. After 2 hours of serum starvation, cells were treated overnight with PAA, PAGly, or PAGln (50 µM), a peroxisome proliferator-activated receptor-alpha (PPAR-α) agonist (GW7647, 1 µM; Cayman Chemical, Cat# 1008613), or vehicle control (DMSO), either alone or in combination with the PPAR-α antagonist (GW6471, 10 µM; Cayman Chemical, Cat# 11697).

### 2.9. RNA-sequencing

#### 2.9.1. RNA isolation

Liver tissue (∼50 mg) or HuH-7 cells were homogenized in TRIzol (1 mL; Qiagen, Cat# 79306). Chloroform (200 µL) was then added, and the samples were mixed and centrifuged to separate phases. The aqueous layer was carefully transferred to a new tube, mixed with 100% isopropanol, vortexed for 15 s, and incubated at −20 °C for at least 1 h. Samples were then centrifuged (20,000 x *g*, 15 min, 4 °C). The resulting RNA pellet was washed twice in 75% ethanol, air-dried, and resuspended in nuclease-free water (Life Technologies, Cat# R0601). RNA concentrations and purity were measured using a NanoDrop (Thermo Fisher Scientific, ND-2000).

#### 2.9.2. Quantitative PCR analysis

Genomic DNA was removed from RNA samples using the DNA-free kit (Invitrogen, Waltham, MA, Cat# AM1906). cDNA was synthesized using the High-Capacity cDNA Reverse Transcription Kit (Thermo Fisher, Waltham, MA, Cat# 4374966). The resulting cDNA (∼100 ng/µL) was mixed with Radiant SYBR Green Master Mix (Alkali Scientific, Cat# QS1020) and quantified by real-time PCR using an Applied Biosystems StepOnePlus thermocycler (Thermo Fisher, Waltham, MA). Fold changes in mRNA expression were calculated using the delta-delta Ct method. The primer list is presented in **Supplementary Table S1**.

#### 2.9.3. Bulk liver RNA-sequencing

mRNA-seq libraries were prepared and sequenced on the Illumina NovaSeq X Plus platform with paired-end 150-bp reads. Approximately 6 Gb of raw data were generated per sample, corresponding to ∼20 million read pairs. After sequencing, paired-end clean reads were aligned to the reference genome using HISAT2 v2.0.5. FeatureCounts v1.5.0-p3 was used to quantify reads mapped to each gene, and fragments per kilobase of transcript values were calculated based on gene length and read counts. Differential expression analysis was performed using the DESeq2 R package v1.20.0. Resulting *P* values were adjusted using the Benjamini-Hochberg method to control the false discovery rate. The clusterProfiler R package was also used to assess enrichment of differentially expressed genes in Kyoto Encyclopedia of Genes and Genomes (KEGG) pathways.

### 2.10. Statistical analysis

Statistical analyses were performed using GraphPad Prism v11. Differences between sexes and experimental groups were evaluated using two-way ANOVA followed by Šídák’s multiple comparisons test. The Kruskal-Wallis test followed by Dunn’s multiple-comparisons test was used for comparisons of multiple groups. Categorical histological score distributions, including NAS scores, were analyzed using Fisher’s exact test. A *P* value < 0.05 was considered statistically significant. Data are presented as mean ± standard error of the mean (SEM).

## 3. Results

### 3.1. PPA exerts sex-dependent effects on body composition and glucose homeostasis

To assess the metabolic impact of phenylalanine-derived microbial metabolites, male and female C57BL/6J mice were fed a high-fat diet and supplemented with PAA or PPA in drinking water for 15 weeks (**Figure 1B**). Both interventions markedly increased circulating downstream metabolites, with PAA supplementation increasing PAGly and PPA supplementation increasing hippuric acid to concentrations comparable to those reported in humans [21, 37]. PAGly levels were higher in females, whereas hippuric acid levels were higher in males (**Figures 1C** and **1D**). PAA also increased circulating PAGln, and both treatments resulted in detectable levels of unmetabolized parent compounds (**Figures S1A-C**).

PPA exerted sex-dependent effects on body composition. In females, PPA increased body mass and adiposity despite food intake being similar to that of control mice. PPA had no effect on body mass or adiposity in males, whereas PAA had no effect in either sex (**Figures 1E-H**).

High-fat feeding impaired glucose tolerance in both sexes, with greater impairment observed in males. Neither PAA nor PPA supplementation altered fasting glucose, insulin, or HOMA-IR (**Figures S2A-C**). However, both metabolites improved glucose tolerance in males, as reflected by reduced glucose excursions during IP-GTT, without altering insulin tolerance. No effects on glucose or insulin tolerance were observed in female mice (**Figure 2**).

**Figure 2.**
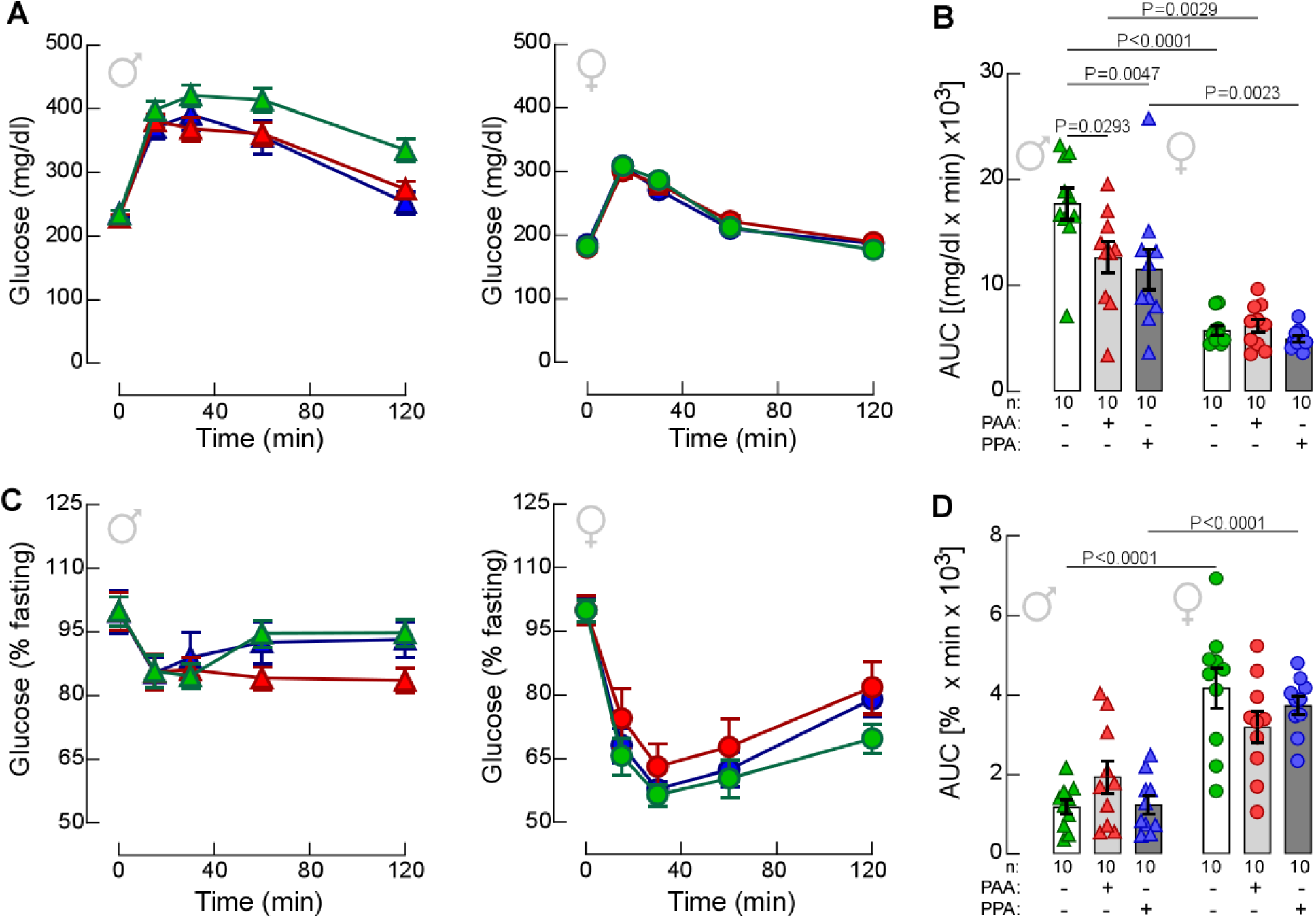
Phenylacetic acid (PAA) and phenylpropionic acid (PPA) improved glucose tolerance in male C57BL/6J mice on a high-fat diet. Blood glucose levels over time following intraperitoneal (**A**) glucose or (**C**) insulin injection in male (left panels) and female (right panels) and (**B** and **D**) corresponding area under the curve (AUC) in male (triangles) and female (circles) mice on a high-fat diet receiving dietary PAA (red) or PPA (blue) supplementation or a control diet (green). Data were analyzed by two-way ANOVA followed by Šídák’s multiple comparisons test. Values are mean ± SEM; n = 10 as indicated.

In summary, these data identify PPA as a sex-dependent regulator of whole-body metabolism, increasing adiposity in females while improving glucose tolerance in males. In contrast, PAA had minimal effects on body composition but selectively improved glucose tolerance in males.

### 3.2. PAA selectively remodels the circulating lipidome

To determine whether PAA and PPA differentially affect systemic lipid metabolism, we performed untargeted lipidomic profiling of plasma. PPA had negligible effects on the plasma lipidome in both sexes (**Figure S3A**). In contrast, PAA induced broad and coordinated changes across multiple lipid classes (**Figure 3A**), with similar directional effects in males and females but a greater overall magnitude of response in females.

**Figure 3.**
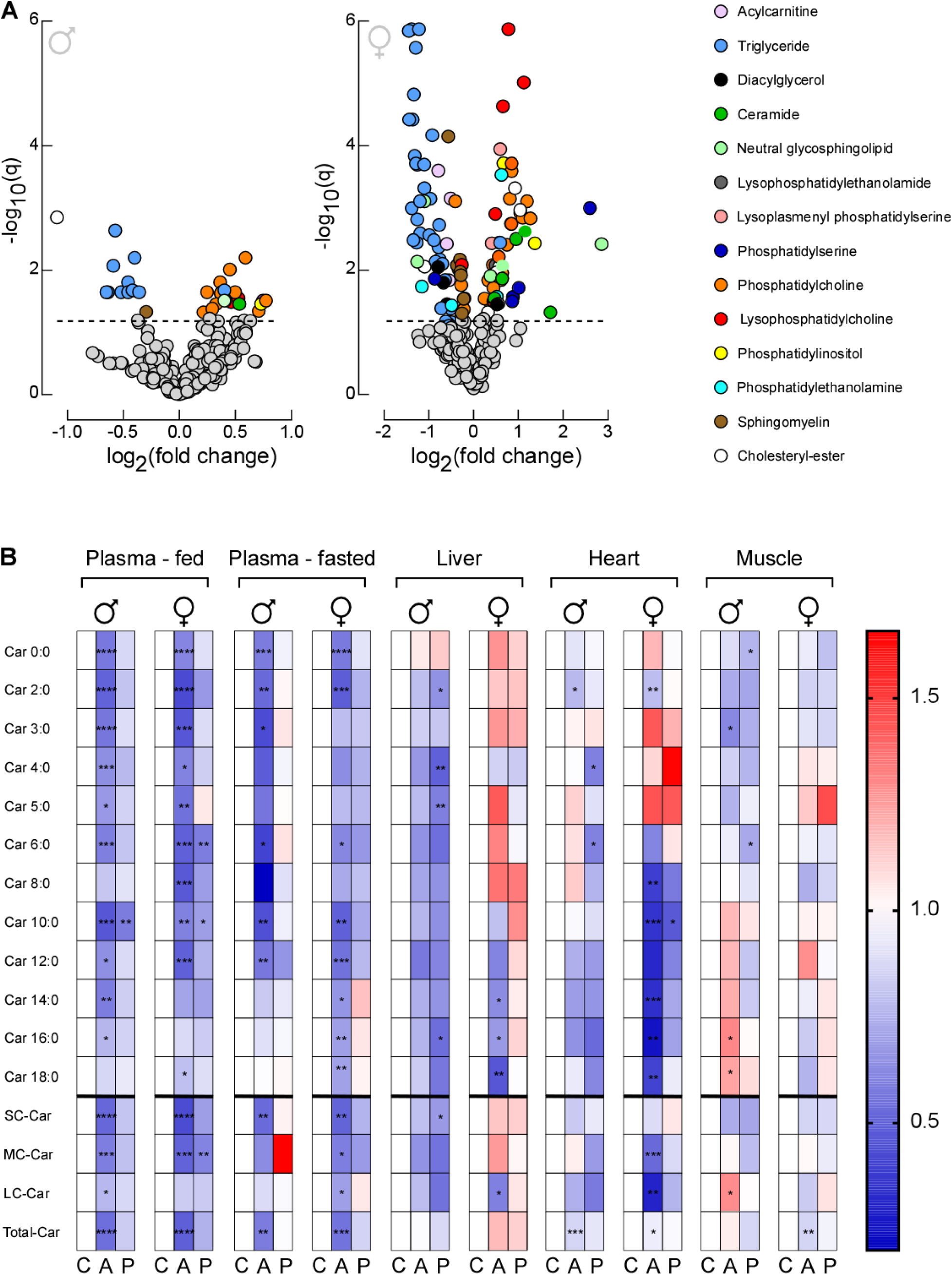
Phenylacetic acid (PAA) alters the circulating lipidome. (**A**) Volcano plots of plasma lipid species in male (left panel) and female (right panel) mice receiving dietary PAA versus controls, showing log-transformed adjusted *P* values and fold changes. Colored circles represent different lipid species, as indicated in the legend. Gray circles represent lipid species with adjusted *P* > 0.05. Negative values indicate downregulated lipids, whereas positive values indicate upregulated lipids following dietary PAA provision. (**B**) Heat map of mean plasma, liver, heart, and muscle levels of free carnitine (Car 0:0) and acylcarnitines (Car 2:0 through Car 18:0), short-chain (SC), medium-chain (MC), and long-chain (LC) acylcarnitines, and the total carnitine pool (Total-Car) in control mice (C) and mice receiving dietary PAA (A) or PPA (P), normalized to the control group. Data were analyzed using the Kruskal-Wallis test followed by Dunn’s multiple-comparisons test (\**P* < 0.05; \*\**P* < 0.01; \*\*\**P* < 0.001; \*\*\*\**P* < 0.0001).

PAA treatment markedly reduced circulating triglycerides, particularly unsaturated triglyceride species, and broadly decreased circulating acylcarnitines. In parallel, phosphatidylcholine and lysophosphatidylcholine species significantly increased. Additional remodeling of sphingolipids, ceramides, phosphatidylserines, and cholesteryl esters was also observed, particularly in female mice (**Figure 3A; Tables S2–S3**). These observations suggest that PAA drives a coordinated reprogramming of systemic lipid metabolism, characterized by reduced circulating neutral lipid and acylcarnitine species together with increased phospholipid abundance.

### 3.3. PAA remodels systemic and tissue carnitine metabolism

Given the prominent reduction in circulating acylcarnitines, we next examined the impact of PAA and PPA on systemic and tissue-specific carnitine metabolism. Targeted metabolomics confirmed a broad reduction in circulating short-, medium-, and long-chain acylcarnitines, as well as free carnitine, in PAA-treated mice (**Figure 3B**). In contrast, PPA had minimal effects on circulating carnitines.

To determine whether increased fecal loss contributed to the reduction in circulating carnitines, fecal carnitine levels were assessed and found to be unchanged or decreased following PAA supplementation, arguing against increased fecal carnitine loss as the cause of its lower circulating pool (**Figure S3B**). We therefore examined tissue-specific carnitine distribution. In the liver, PAA treatment did not alter total carnitine content in male mice. In female mice, PAA slightly shifted hepatic carnitine composition, characterized by reduced long-chain acylcarnitines while total hepatic carnitine content remained preserved. In the heart, PAA selectively reduced medium- and long-chain acylcarnitines in females, while having minimal effects in males. Skeletal muscle was comparatively unaffected, although male mice exhibited increased C16 and C18 acylcarnitines and females showed a modest reduction in total carnitine pool (**Figure 3B**).

Notably, circulating free fatty acid levels were unchanged (**Figure S3C**), indicating that these changes occurred without detectable alterations in steady-state circulating free fatty acid concentrations. Together with the observed reduction in circulating triglycerides (**Figure 3A**), the marked contraction of the circulating carnitine pool contrasted with relatively modest and tissue-specific changes in tissue acylcarnitine composition, suggesting that PAA alters systemic handling of lipids and carnitine-associated metabolites.

### 3.4. PAA-induced lipid remodeling occurs without steatosis progression or overt liver injury

Despite these profound metabolic changes, PAA did not significantly increase hepatic lipid accumulation. Total liver triglyceride content was elevated only in PAA-treated male mice, while hepatic cholesterol and bile acid levels were largely unchanged (**Figures 4A-C; Figure S4A-B**). Histological analysis revealed no significant differences in steatosis severity or NAS between treatment groups (**Figures 4D-E; Figure S4C**).

**Figure 4.**
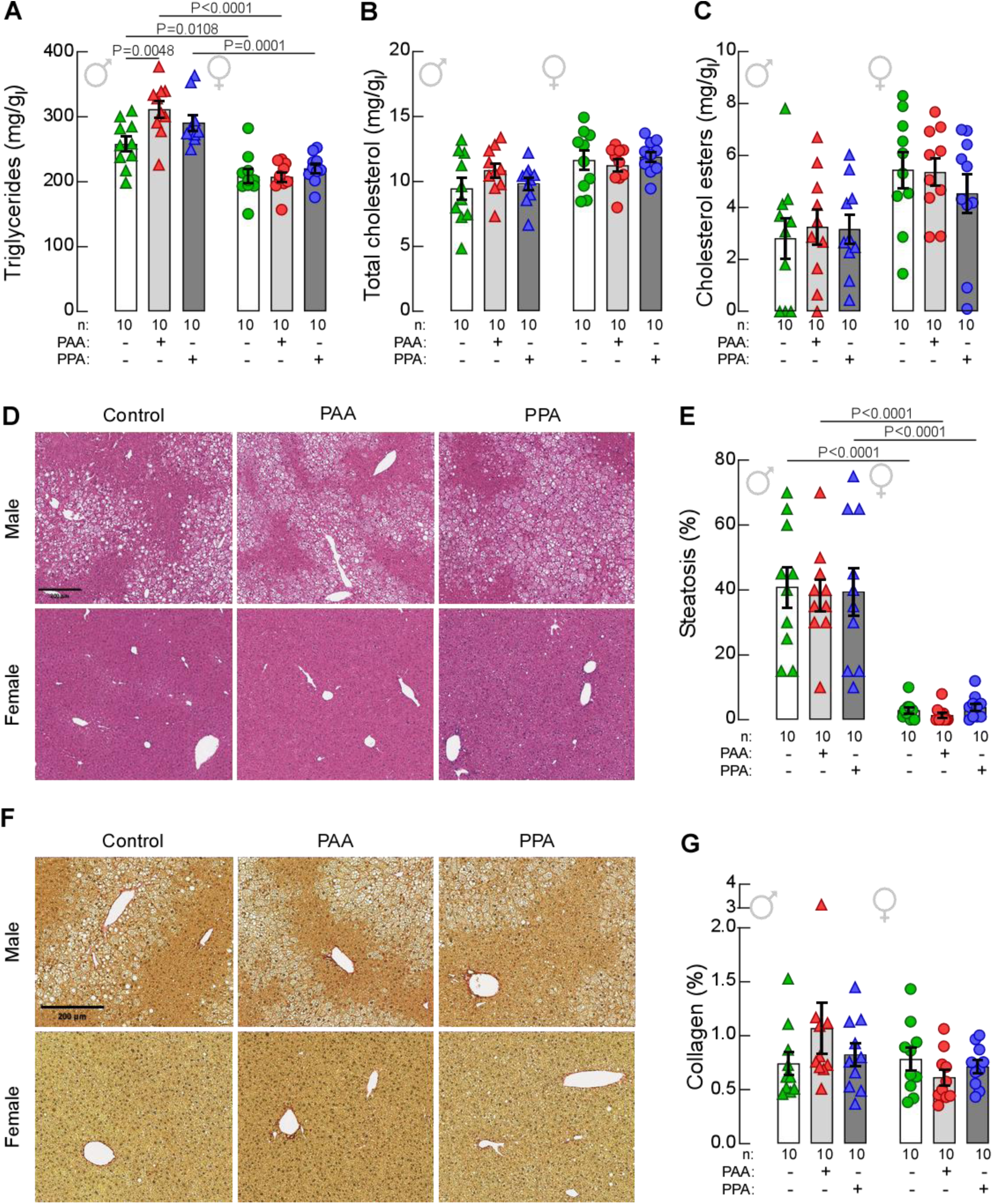
Dietary provision of phenylacetic acid (PAA) and phenylpropionic acid (PPA) do not affect liver steatosis in a mouse model of diet-induced obesity. Liver (**A**) triglycerides, (**B**) total cholesterol, and (**C**) cholesteryl esters. (**D**) Representative H&E-stained liver sections; scale bar = 200 µm. (**E**) Percentage of liver steatosis. (**F**) Representative Picrosirius red-stained liver sections; scale bar = 200 µm. (**G**) Percentage of collagen in male (triangles) and female (circles) mice on a high-fat diet receiving dietary PAA (red) or PPA (blue) supplementation or a control diet (green). Data were analyzed using two-way ANOVA followed by Šídák’s multiple-comparisons test. Values are mean ± SEM; n = 10, as indicated.

PAA treatment induced hepatomegaly selectively in female mice, as evidenced by increased liver mass (**Figures S4D-E**); however, it was not accompanied by elevated circulating ALT, AST, or ALP activity, indicating the absence of overt hepatocellular injury. PPA treatment also did not increase ALT, AST, or ALP activities, consistent with minimal lipidomic changes (**Figures S4F-H**).

Histological analysis revealed mild inflammatory infiltrates in treated females but no evidence of hepatocellular ballooning, steatohepatitis, or fibrosis. Consistent with these findings, the expression of pro-inflammatory markers, such as TNF-α, was not significantly increased (**Figures 4F-G; Figures S4I-J**). In summary, PAA induced adaptive hepatic remodeling, likely reflecting increased metabolic capacity. There was no evidence of hepatocellular stress following PPA treatment.

### 3.5. Hepatic transcriptomic response suggests PAA-driven remodeling of lipid handling, peroxisomal fatty acid metabolism, and xenobiotic pathways

To further define the hepatic response to phenolic acid supplementation, we performed bulk RNA-sequencing of liver tissue from male and female mice. PAA induced substantial transcriptional remodeling, with markedly stronger effects in females than males (**Figure 5A, Tables S4-S5**). In males, the most prominently induced genes included members of the *Cyp4a* family, as well as *Ehhadh, Acot1, Dusp8,* and *Retsat*, implicating pathways involved in fatty acid metabolism, ω-hydroxylation, peroxisomal lipid processing, and metabolic adaptation. Female mice exhibited a broader response characterized by induction of genes involved in lipid uptake, storage, mobilization, and fatty acid catabolism, including *Acox1, Acaa1b, Ucp3, Mogat1, Aqp7, Cidec*, and *Cfd*. KEGG pathway analysis identified enrichment of PPAR signaling, fatty acid metabolism, fatty acid degradation, fatty acid elongation, and retinol metabolism, with a similar but more extensive pattern observed in females (**Figure 5B**).

**Figure 5.**
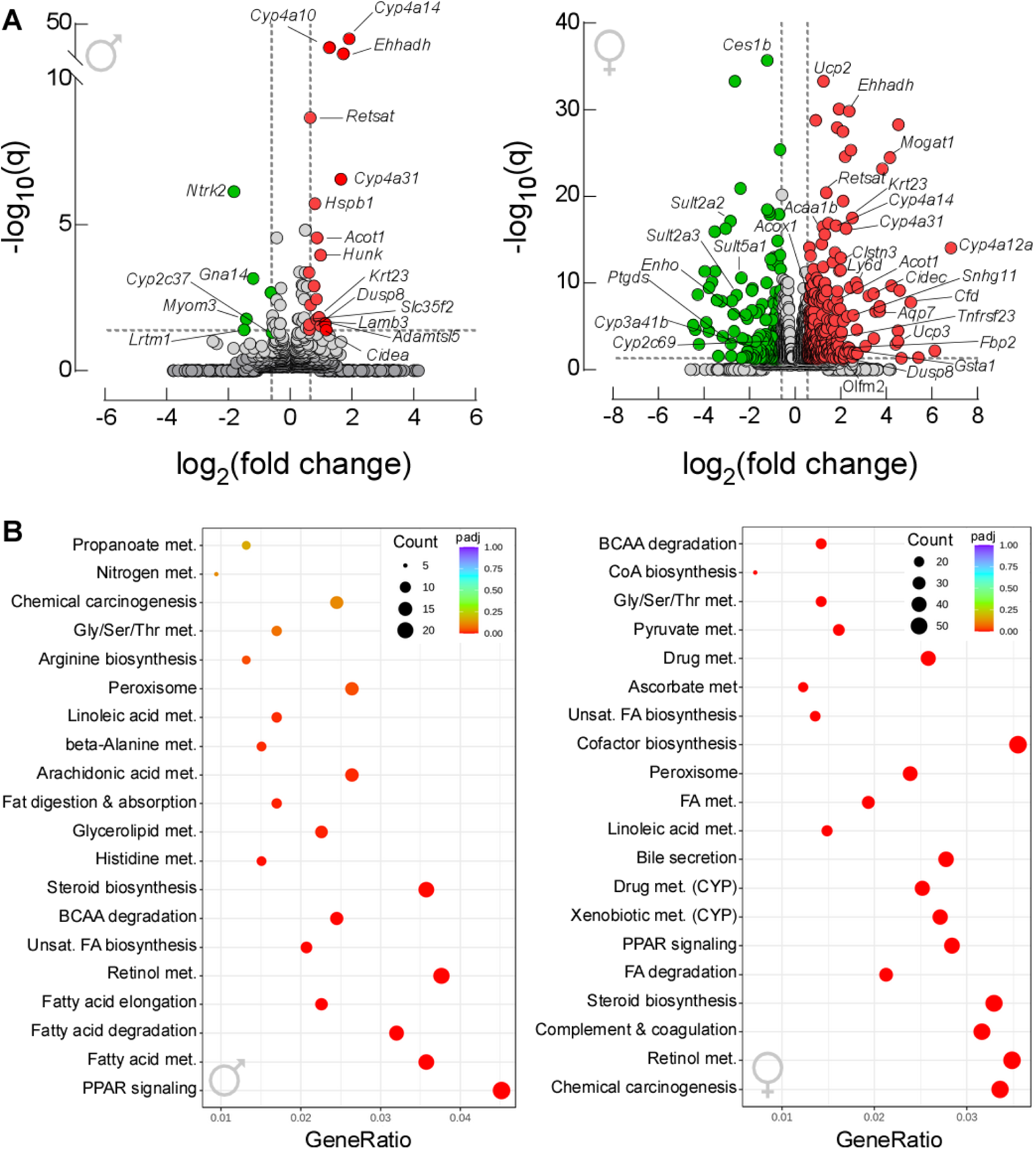
Phenylacetic acid (PAA) dysregulates genes associated with fatty acid metabolism and lipid homeostasis. (**A**) Volcano plots of hepatic gene expression in male (left panel) and female (right panel) mice receiving dietary PAA versus controls, showing log-transformed adjusted *P* values and fold changes. Green circles represent genes with significantly higher expression in the control group, and red circles represent genes upregulated following dietary PAA provision. Gray circles represent genes with adjusted *P* > 0.05 and/or │log2 fold change│< 0.58. (**B**) KEGG dot plots showing the top 20 pathways dysregulated by dietary PAA provision in male (left panel) and female (right panel) mice.

Several genes previously associated with steatosis, liver injury, or carcinogenesis were also altered following PAA treatment. However, these transcriptional changes occurred in the absence of corresponding histologic or biochemical evidence of liver pathology. Although *Col1a1* expression was modestly increased in females, other profibrotic markers, including *Acta2*, *Col3a1*, and fibrogenic growth factors, were unchanged, arguing against activation of a broader fibrogenic program (**Tables S4-S5**).

These findings suggest that PAA remodels hepatic lipid-processing pathways. The coordinated induction of genes involved in PPAR signaling, ω-oxidation (*Cyp4a*), and peroxisomal fatty acid metabolism (*Acox1*, *Ehhadh*, *Acaa1b*) is consistent with activation of transcriptional programs involved in fatty acid processing and acyl-group metabolism rather than increased hepatic lipid storage. Importantly, these molecular changes occurred without worsening steatosis, liver injury, or inflammation, indicating an adaptive hepatic remodeling response rather than overt hepatotoxicity.

In contrast, PPA exerted minimal effects on hepatic gene expression, liver lipid composition, and markers of liver injury (**Figure S5, Tables S6-S7**). Collectively, these data demonstrate that PAA, but not PPA, induces a sex-dependent hepatic metabolic remodeling program characterized by activation of lipid-handling, peroxisomal, and xenobiotic pathways, accompanied by reduced circulating triglycerides and acylcarnitines, while preserving overall liver health.

### 3.6. PAA promotes PPAR-α-responsive gene expression in hepatocytes

Liver RNA-sequencing identified enrichment of PPAR signaling pathways and induction of multiple established PPAR-α target genes following PAA treatment. To determine whether PAA and its downstream host metabolites directly regulate PPARα-responsive transcriptional programs, HuH-7 hepatocytes were treated with PAA, PAGly, or PAGln. All three metabolites increased expression of the canonical PPAR-α target gene *ACOX1* (**Figure 6A**). Pharmacological inhibition of PPAR-α abolished PAA-induced *ACOX1* expression, supporting a role for PPAR-α signaling in mediating the transcriptional response to PAA (**Figure 6B**). The antagonist also attenuated *ACOX1* induction by PAGly and PAGln, although these effects did not reach statistical significance. Together, these findings support involvement of PPAR-α-responsive transcriptional pathways in the hepatic response to PAA and are consistent with the transcriptional remodeling observed *in vivo*.

**Figure 6.**
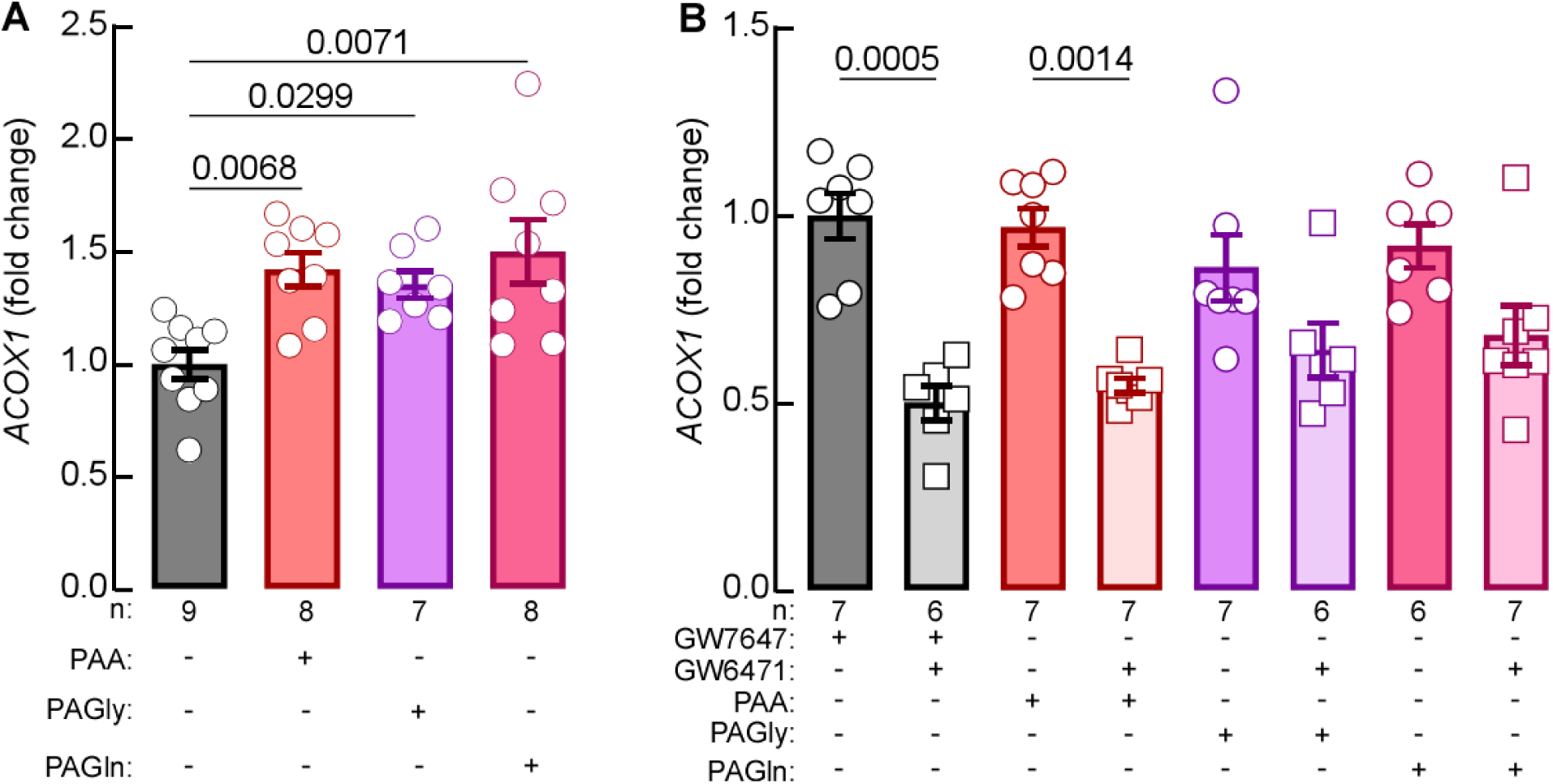
Phenylacetic acid (PAA) and its downstream host metabolites induce PPAR-α-responsive gene expression in HuH-7 hepatocytes. (**A**) Fold change in *ACOX1* expression in HuH-7 cells treated with PAA, phenylacetylglycine (PAGly), phenylacetylglutamine (PAGln), or vehicle control (DMSO). (**B**) Fold change in *ACOX1* expression in HuH-7 cells treated with the PPAR-α agonist GW7647, PAA, PAGly, or PAGln, with or without the PPAR-α antagonist GW6471. Data were analyzed using the Kruskal-Wallis test followed by Dunn’s multiple-comparisons test. Values are mean ± SEM; n = 6–9, as indicated, from two independent experiments.

## 4. Discussion

Given the strong associations between gut microbial composition and host metabolic diseases, targeting the microbiome has emerged as a promising therapeutic strategy. However, directly manipulating complex microbial communities remains challenging. An alternative and potentially more tractable approach is to target microbiome-derived metabolites, which represent discrete, chemically defined mediators of host–microbe interactions. Achieving this goal requires an understanding of how individual metabolites regulate host physiology beyond observational clinical correlations. Here, we demonstrate that two structurally related phenylalanine-derived microbial metabolites, PAA and PPA, exert fundamentally distinct and sex-dependent effects on host metabolism, revealing divergent metabolic axes through which the microbiome can influence metabolic health.

PAA and its downstream host metabolite PAGln have been consistently associated with obesity, insulin resistance, and T2DM in human studies [26, 28, 40–43]. Surprisingly, despite these associations, chronic dietary provision of PAA in a mouse model of diet-induced obesity did not affect body mass, adiposity, fasting glucose, or insulin sensitivity. Instead, PAA induced a profound reprogramming of hepatic metabolism characterized by coordinated changes in lipidomic profiles, transcriptional pathways, and metabolite profiles. Specifically, PAA reduced circulating unsaturated triglycerides as well as free and acylcarnitines in plasma. These changes were accompanied by increased phosphatidylcholine and lysophosphatidylcholine species, consistent with enhanced phospholipid remodeling and altered lipid partitioning.

Further, the marked reduction in circulating carnitines was not accompanied by evidence of global tissue carnitine depletion. Total hepatic carnitine content remained unchanged, and fecal carnitine levels were not increased. Instead, PAA produced sex- and tissue-specific remodeling of acylcarnitine pools, with reductions in hepatic long-chain acylcarnitines and cardiac medium-and long-chain acylcarnitines observed primarily in females. These findings indicate redistribution of carnitine-associated metabolites across tissues, although the underlying metabolic fluxes cannot be determined from steady-state acylcarnitine measurements alone.

The coordinated induction of genes involved in PPAR signaling, ω-hydroxylation (*Cyp4a* family), and peroxisomal lipid metabolism (*Acox1*, *Ehhadh*, *Acaa1b*) is consistent with activation of hepatic lipid-processing pathways. Because transcriptomic and steady-state metabolomic measurements do not directly assess metabolic flux, the extent to which these changes reflect altered fatty acid metabolism or acyl-group handling remains to be determined. The broader lipidomic, acylcarnitine, and hepatic transcriptional responses observed in females suggest that sex is an important determinant of host responsiveness to PAA.

Consistent with the hepatic transcriptomic findings, PAA, PAGly, and PAGln increased expression of the PPAR-α target gene *ACOX1* in HuH-7 cells. Pharmacologic inhibition of PPAR-α abolished the response to PAA, supporting involvement of PPAR-α-responsive transcriptional pathways in the hepatic response to PAA.

Despite this extensive metabolic reprogramming, PAA did not increase hepatic lipid accumulation or induce overt liver injury. Total liver triglyceride content and histological steatosis were largely unchanged, and circulating ALT, AST, and ALP levels remained within the normal range. Although female mice developed hepatomegaly and exhibited modest changes in expression of selected genes previously linked to liver disease or fibrosis, these changes occurred in the absence of histological evidence of steatohepatitis, fibrosis, or significant hepatic inflammation. Thus, the transcriptional and metabolic effects of PAA appear to reflect adaptive remodeling of hepatic lipid metabolism rather than hepatotoxicity. The absence of substantial changes in liver fat content despite marked alterations in lipid metabolic pathways further suggests that PAA primarily affects lipid flux and processing rather than steady-state hepatic lipid storage. Interestingly, our findings differ from a recent report demonstrating that PAA-derived lysine phenylacetylation impairs mitochondrial function and promotes features of MASLD [44]. One possible explanation for these apparently divergent findings is that PAA engages both adaptive and maladaptive pathways, with the balance between PPAR-α-associated metabolic remodeling and phenylacetylation-dependent mitochondrial stress being influenced by disease state, exposure level, sex, and host metabolic context. Further studies will be required to determine how these pathways interact and whether PPAR-α activation serves as a compensatory response to increased PAA exposure.

In contrast to PAA, PPA had minimal impact on hepatic lipid metabolism, transcriptomic programs, or circulating lipid profiles. Nevertheless, PPA induced sex-dependent physiological effects. In female mice, PPA promoted weight gain and adiposity despite reduced food intake, suggesting altered energy partitioning toward lipid storage. This was without substantial changes in hepatic gene expression or lipid accumulation, indicating a disconnect between systemic phenotype and hepatic metabolic remodeling. In male mice, PPA improved glucose tolerance without affecting body mass or adiposity, consistent with prior studies demonstrating beneficial effects of hippuric acid on glucose metabolism [31]. These findings reveal that PPA primarily affects systemic metabolic regulation rather than hepatic metabolic pathways.

The marked sexual dimorphism observed in response to both metabolites highlights the importance of host context in determining the physiological effects of microbiome-derived metabolites. Most prior studies examining phenylalanine-derived metabolites have been conducted predominantly in male animals, potentially overlooking important sex-specific responses. Our findings demonstrate that the metabolic consequences of these metabolites depend not only on metabolite structure but also on the physiological environment in which they act and challenge the assumption that microbially derived metabolites exert uniform or predictable effects based solely on their clinical associations. Although PAA and PPA originate from the same precursor, they engage distinct biological processes and produce markedly different host responses. PAA acts primarily as a modulator of hepatic lipid metabolism, promoting PPAR-α-associated remodeling of fatty acid processing, peroxisomal metabolism, and systemic and tissue carnitine metabolism without inducing overt liver injury. In contrast, PPA functions principally as a modulator of whole-body metabolic physiology, exerting sex-dependent effects on adiposity and glucose homeostasis largely independent of major hepatic metabolic remodeling.

## 5. Conclusion

In conclusion, our study demonstrates that targeting individual microbial metabolites reveals a previously unappreciated level of specificity in microbiome–host interactions. Rather than acting as generalized mediators of metabolic health or disease, phenylalanine-derived metabolites selectively regulate distinct metabolic pathways and physiological processes. These findings highlight the need for mechanistic dissection of individual metabolite pathways and suggest that effective microbiome-based therapies will require precise targeting of metabolite-specific functions rather than broad manipulation of microbial communities.

## Ethics statement

All animal studies were approved by the Cleveland Clinic Research Institutional Animal Care and Use Committee.

## Data availability

Untargeted lipidomics data generated in this study are available on Zenodo (DOI: 10.5281/zenodo.18166719), and raw bulk liver RNA-sequencing data are deposited in the NCBI Sequence Read Archive under accession number PRJNA1422046. Any additional information required to reanalyze the data reported in this paper is available from the lead contact upon reasonable request.

## ACKNOWLEDGMENTS

The graphical abstract and Figure 1B were created using BioRender.com. We thank Apryl Helmick and Ajay Zalavadia of the Cleveland Clinic Research Imaging Core for performing the histological staining of liver tissues and the quantitative analysis of liver fibrosis, respectively.

## Abbreviations

ALP: alkaline phosphatas
ALT: alanine aminotransferase
ANN: artificial neural network
ANOVA: analysis of variance
AST: aspartate aminotransferase
AUC: area under the curve
BA: bile acid
Car: carnitine
DMSO: dimethyl sulfoxide
FXR: farnesoid X receptor
H&E: hematoxylin and eosin
HFD: high-fat diet
HOMA-IR: homeostatic model assessment for insulin resistance
IP: intraperitoneal
IP-GTT: intraperitoneal glucose tolerance test
IP-ITT: intraperitoneal insulin tolerance test
KEGG: Kyoto Encyclopedia of Genes and Genomes
LC: long-chain
LC-MS/MS: liquid chromatography–tandem mass spectrometry
MASLD: metabolic dysfunction-associated steatotic liver disease
MC: medium-chain
MRM: multiple reaction monitoring
MS/MS: tandem mass spectrometry
m/z: mass-to-charge ratio
NAS: nonalcoholic fatty liver disease activity score
PAA: phenylacetic acid
PAGln: phenylacetylglutamine
PAGly: phenylacetylglycine
PPA: phenylpropionic acid
PPAR-α: peroxisome proliferator-activated receptor alpha
SC: short-chain
SCFA: short-chain fatty acid
SEM: standard error of the mean
T2DM: type 2 diabetes mellitus
TGR5: Takeda G protein-coupled receptor 5

