## Supplementary Files for "Gut microbially produced phenolic acids have differential and sex-dependent effects on host metabolic health"

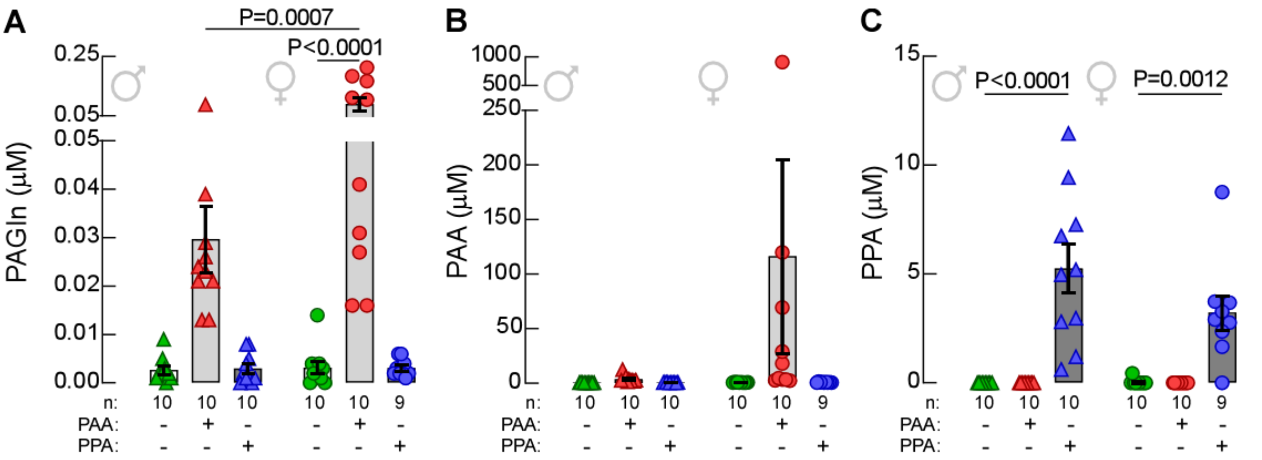


**Supplementary Figure S1.** Plasma levels of (**A**) phenylacetylglutamine (PAGln), (**B**) phenylacetic acid (PAA), and (**C**) phenylpropionic acid (PPA) in male (triangles) and female (circles) mice on a high-fat diet receiving dietary PAA (red) or PPA (blue) supplementation or a control diet (green). Data were analyzed by two-way ANOVA followed by Šídák's multiple comparisons test. Values are mean ± SEM; n = 9-10 as indicated.

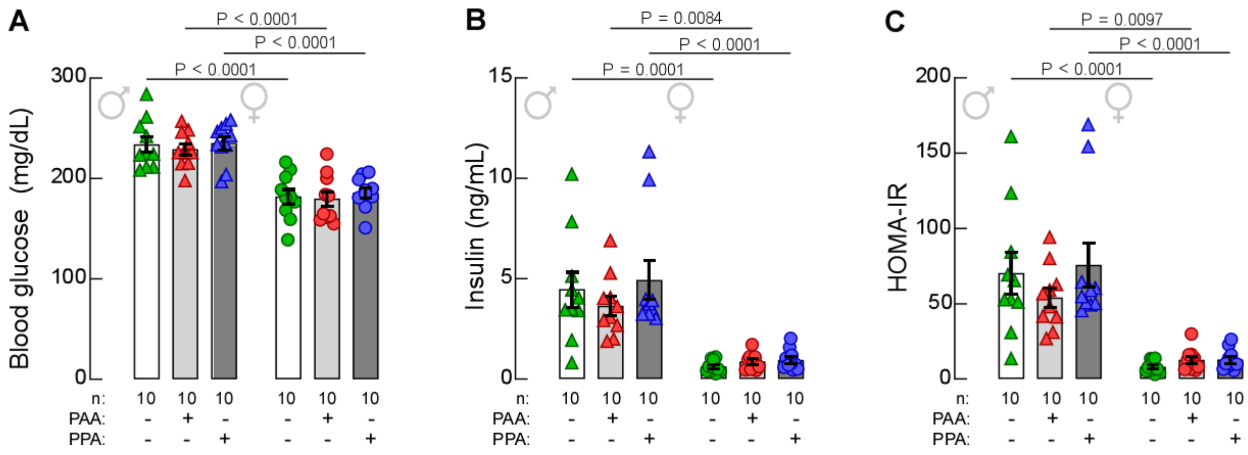


**Supplementary Figure S2.** Fasting (**A**) blood glucose levels, (**B**) insulin levels, and (**C**) HOMA-IR in male (triangles) and female (circles) mice on a high-fat diet receiving dietary PAA (red) or PPA (blue) supplementation or a control diet (green). Data were analyzed by two-way ANOVA followed by Šídák's multiple comparisons test. Values are mean ± SEM; n = 10 as indicated.

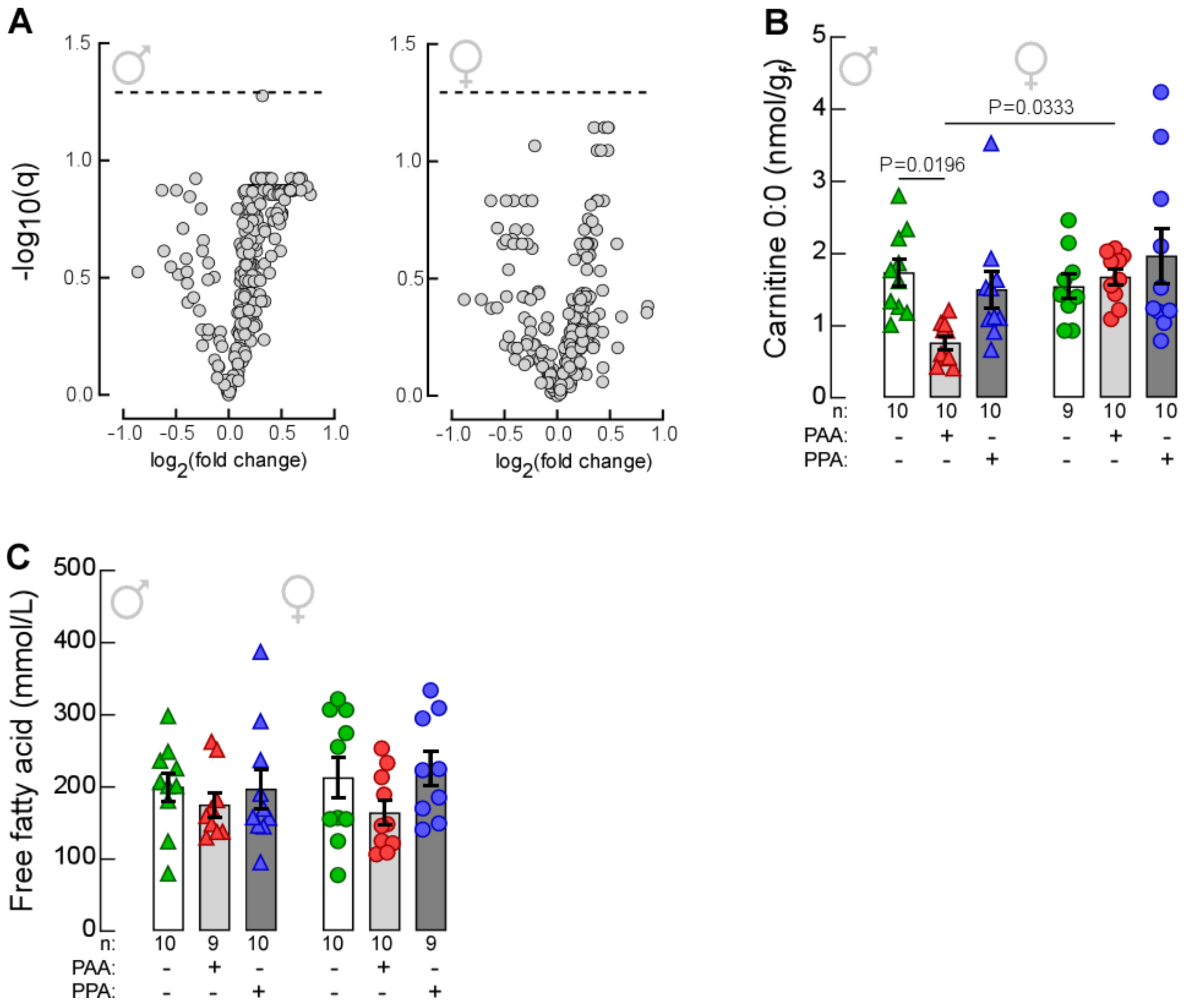


**Supplementary Figure S3.** (**A**) Volcano plot of plasma lipid species in male (left panel) and female (right panel) mice with dietary provision of PPA versus control. Gray circles represent lipid species with adjusted $P > 0.05$. (**B**) Fecal levels of carnitine (Car 0:0) and (**C**) circulating total free fatty acids in male (triangles) and female (circles) mice on a high-fat diet receiving dietary PAA (red) or PPA (blue) supplementation or a control diet (green). Data were analyzed by two-way ANOVA followed by Šídák's multiple comparisons test. Values are mean ± SEM; n = 9-10 as indicated.

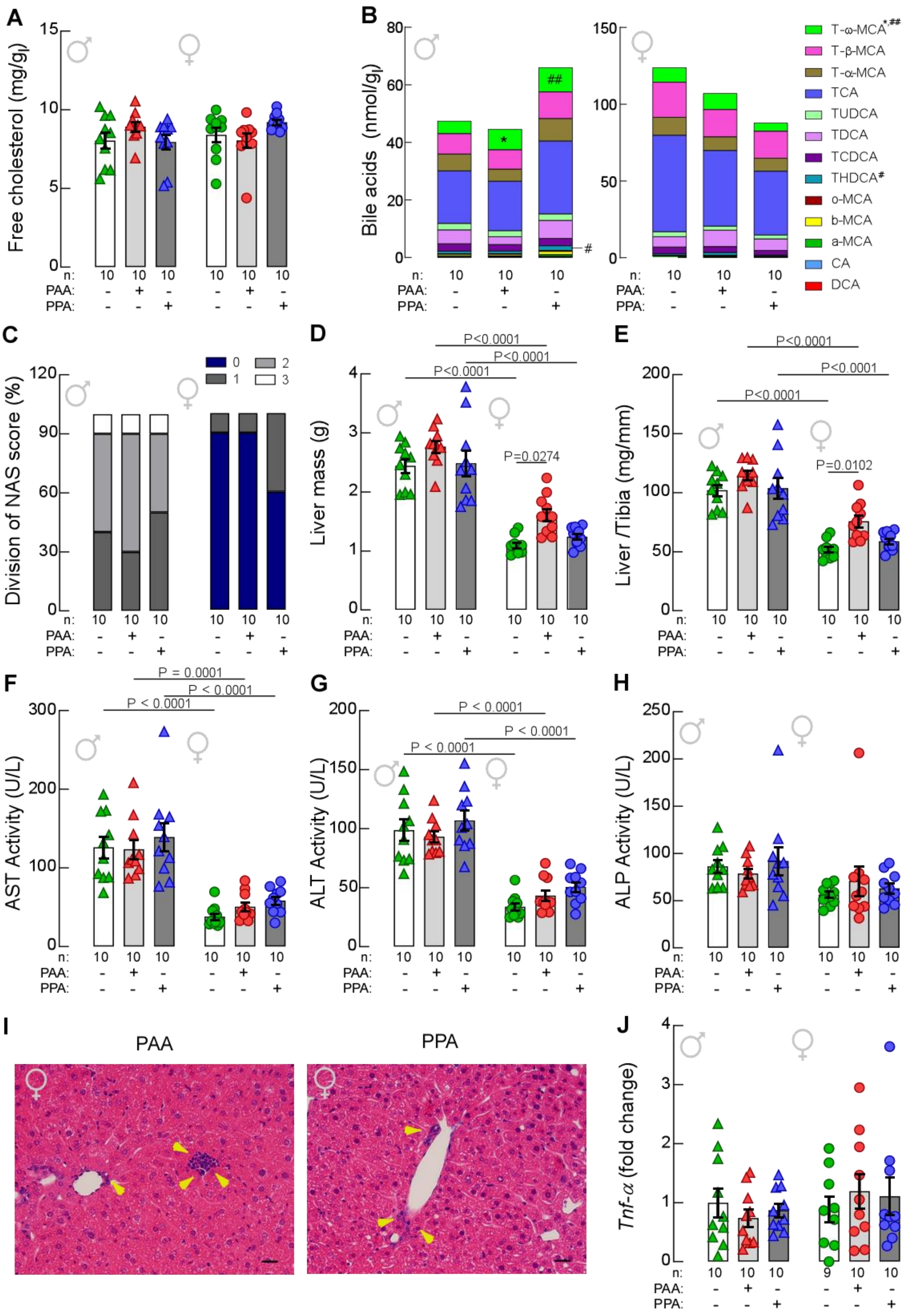

A
Free cholesterol (mg/gl)
B
Bile acids (nmol/gl)
T-ω-MCA*,##
T-β-MCA
T-α-MCA
TCA
TUDCA
TDCA
TCDCA
THDCA#
o-MCA
b-MCA
a-MCA
CA
DCA
C
Division of NAS score (%)
0
1
2
3
D
Liver mass (g)
P<0.0001
P<0.0001
P<0.0001
P=0.0274
E
Liver /Tibia (mg/mm)
P<0.0001
P<0.0001
P<0.0001
P=0.0102
F
AST Activity (U/L)
P = 0.0001
P < 0.0001
P < 0.0001
G
ALT Activity (U/L)
P < 0.0001
P < 0.0001
P < 0.0001
H
ALP Activity (U/L)
I
PAA
PPA
J
Tnf-α (fold change)
n:
PAA:
PPA:

**Supplementary Figure S4.** (**A**) Free liver cholesterol normalized to liver mass ($g_l$) in male (triangles) and female (circles) mice on a high-fat diet receiving dietary PAA (red) or PPA (blue) supplementation or a control diet (green). (**B**) Levels of hepatic bile acids in male (left panel) and female (right panel) mice after dietary provision of PAA and PPA; individual bile acids are indicated in the legend on the right. (**C**) Distribution of liver NAS scores across treatment groups. (**D**) Liver mass and (**E**) liver mass normalized to tibia length. Plasma (**F**) aspartate aminotransferase (AST), (**G**) alanine aminotransferase (ALT), and (**H**) alkaline phosphatase (ALP) activities. (**I**) Representative H&E-stained liver sections of PAA-treated (left panel) and PPA-treated (right panel) female mice. Arrowheads point to focal lymphocytes. Scale bar = 20 µm. (**J**) Hepatic *Tnf-α* mRNA expression in male (triangles) and female (circles) mice on a high-fat diet with and without dietary provision of PAA (red) and PPA (blue) versus control (green). (**A, D-H**, and **J**) Data were analyzed by two-way ANOVA followed by Šídák's multiple comparisons. Values are represented as mean ± SEM; n = 9-10 as indicated. For panel **B**, data were analyzed using the Kruskal-Wallis test followed by Dunn's multiple comparisons test. In male mice, bile acids that differed significantly from control are marked as follows: $^{*}P < 0.05$ for the PAA group; $^{\#}P < 0.05$ and $^{\#\#}P < 0.01$ for the PPA group. (**C**) Data were analyzed using Fisher's exact test.

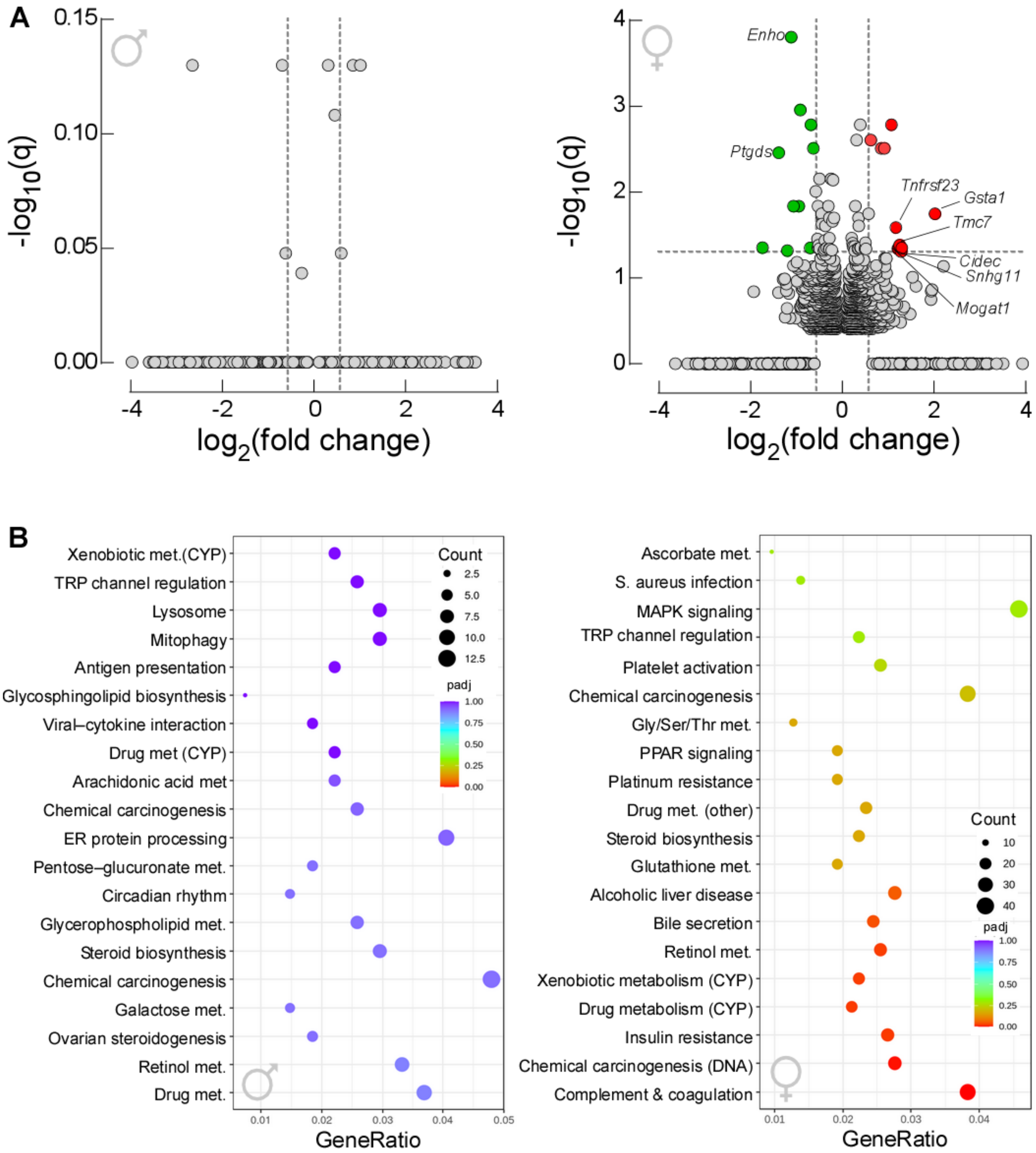


**Supplementary Figure S5.** (**A**) Volcano plot of liver expressed genes in male (left panel) and female (right panel) mice with dietary provision of PPA versus control. Green circles represent genes significantly higher in the control group and blue circles represent genes upregulated with dietary provision of PPA. Gray circles represent genes with adjusted *P* > 0.05 and/or $|\log_2$ fold change$|$ < 0.58. (**B**) KEGG analysis dot plots for the top 20 pathways altered by dietary PPA supplementation in male (left panel) and female (right panel) mice.
